# The Lid/KDM5 histone demethylase complex activates a critical effector of the oocyte-to-zygote transition

**DOI:** 10.1101/682468

**Authors:** Daniela Torres-Campana, Shuhei Kimura, Guillermo A. Orsi, Béatrice Horard, Gérard Benoit, Benjamin Loppin

## Abstract

Following fertilization of a mature oocyte, the formation of a diploid zygote involves a series of coordinated cellular events that ends with the first embryonic mitosis. In animals, this complex developmental transition is almost entirely controlled by maternal gene products. How such a crucial transcriptional program is established during oogenesis remains poorly understood. Here, we have performed an shRNA-based genetic screen in *Drosophila* to identify genes required to form a diploid zygote. We found that the Lid/KDM5 histone demethylase and its partner, the Sin3A-HDAC1 deacetylase complex, are necessary for sperm nuclear decompaction and karyogamy. Surprisingly, transcriptomic analyses revealed that these histone modifiers are required for the massive transcriptional activation of *deadhead* (*dhd*), which encodes a maternal thioredoxin involved in sperm chromatin remodeling. Unexpectedly, while *lid* knock-down tends to slightly favor the accumulation of its target, H3K4me3, on the genome, this mark was lost at the *dhd* locus. We propose that Lid/KDM5 and Sin3A cooperate to establish a local chromatin environment facilitating the unusually high expression of *dhd*, a key effector of the oocyte-to-zygote transition.

## Introduction

In obligate sexually reproducing animals, fertilization allows the formation of a diploid zygote through the association of two haploid gametes of highly different origins and structures. Generally, the spermatozoon delivers its compact nucleus within the egg cytoplasm, along with a pair of centrioles, while the egg provides one haploid meiotic product and all resources to sustain zygote formation [1]. In some species, this maternal control extends to early embryo development, as in *Drosophila melanogaster*, where the initial amplification of embryo cleavage nuclei occurs without significant zygotic transcription [2]. The developmental potential of the egg is thus initially dependent on the establishment of a highly complex transcriptional program in female germ cells. During *Drosophila* oogenesis, the bulk of transcriptional activity takes place in the fifteen interconnected large polyploid germline nurse cells that deposit gene products and metabolites in the cytoplasm of the growing oocyte [3].

One of the earliest events of the oocyte-to-zygote transition is the rapid transformation of the fertilizing sperm nucleus into a functional male pronucleus. In *Drosophila*, the needle-shaped, highly compact sperm nucleus is indeed almost entirely organized with non-histone, Sperm Nuclear Basic Proteins (SNBPs) of the protamine-like type [4]. Male pronucleus formation begins with the genome-wide replacement of SNBPs with maternally supplied histones, a process called sperm chromatin remodeling, which is followed by extensive pronuclear decondensation [1]. Finally, zygote formation involves the coordinated migration and apposition of male and female pronuclei and the switch from meiotic to mitotic division within the same cytoplasm.

Here, we report the results of a genetic screen specifically designed to find new genes required for the oocyte-to-zygote transition in *Drosophila*. Our screen identified two histone modifiers, the Lid/KDM5 histone H3K4me3 demethylase and the Sin3A-HDAC1 histone deacetylase complex, which are both required for the integration of paternal chromosomes into the zygote. These interacting epigenetic factors are known to regulate the expression of hundreds of genes in somatic tissues but their role in the establishment of the ovarian transcriptome is unknown. Strikingly, RNA-Sequencing analyses revealed that, despite the modest impact of their depletion on ovarian transcripts, Lid and Sin3A are critically required for the massive expression of *deadhead* (*dhd*), a key effector of the oocyte-to-zygote transition [5,6]. Furthermore, we demonstrate that germline knock-down of these histone modifiers specifically prevent sperm chromatin remodeling through a mechanism that depends on the DHD thioredoxin.

## Results & Discussion

### A maternal germline genetic screen for gTBLhaploid embryo development

We performed an *in vivo* RNA interference screen in the female germline to identify genes required for the integration of paternal chromosomes in the zygote. In *Drosophila*, failure to form a male pronucleus following fertilization is generally associated with the development of haploid embryos that possess only maternally-derived chromosomes (gynohaploid embryos) and that never hatch [7]. We chose to screen transgenic lines from the TRiP collection that express small hairpin RNAs (shRNAs) under the control of the Gal4 activator [8]. We selected shRNA lines that targeted genes with known or predicted chromatin-related function and that show adult ovarian expression (Flybase). Among the 374 tested TRiP lines, 157 (41.9 %) induced female sterility or severely reduced fertility when induced with the germline-specific *P{GAL4::VP16-nos.UTR}*^*MVD1*^ Gal4 driver (*nos*-Gal4), thus underlying the importance of chromatin regulation for oogenesis and early embryo development (**Table EV1**). We then specifically searched for shRNAs that induced a maternal effect embryonic lethal phenotype associated with gynohaploid development (Fig. 1A). Gynohaploid embryos can be efficiently identified by scoring the zygotic expression of a paternally-transmitted *P[g-GFP::cid]* transgene at the gastrulation stage or beyond [9]. Among the sterile lines with late developing embryos (class 4 in Fig. 1A and **Table EV1**) that were identified, four shRNA lines (GLV21071, GL00612, HMS00359 and HMS00607) produced embryos that were negative for GFP::CID (Fig. 1B). Note that none of these shRNAs induced complete female sterility and about 1 to 4% of embryos hatched and were thus diploid (Table 1).

**Table 1.** Embryo hatching rates

| Female genotype | Male genotype | Number of eggs | Hatch. rate (%) |
| --- | --- | --- | --- |
| *w<sup>1118</sup>* | *w<sup>1118</sup>* | 344 | 97.67 |
| *dhd<sup>J5</sup>* | *w<sup>1118</sup>* | 375 | 0.00 |
| *dhd<sup>J5</sup>/FM7c* | *w<sup>1118</sup>* | 573 | 78.36 |
| *MTD>+* | *w<sup>1118</sup>* | 1561 | 98.27 |
| *MTD>sh-lid22* | *w<sup>1118</sup>* | 1403 | 2.14 |
| *MTD>sh-lid21* | *w<sup>1118</sup>* | 1144 | 1.05 |
| *MTD>sh-Sin3A* | *w<sup>1118</sup>* | 1221 | 0.25 |
| *MTD>sh-rpd3* | *w<sup>1118</sup>* | 521 | 4.03 |
| *MTD>sh-Hira* | *w<sup>1118</sup>* | 315 | 0.00 |
| **Rescue with *P[dhd<sup>WT</sup>]* transgene** |  |  |  |
| *dhd<sup>J5</sup> ; P[dhd]* | *w<sup>1118</sup>* | 663 | 85.67 |
| *dhd<sup>J5</sup> ; P[dhd]/TM2* | *w<sup>1118</sup>* | 884 | 94.57 |
| *MTD>sh-lid22 , P[dhd]* | *w<sup>1118</sup>* | 1159 | 4.57 |
| **Rescue with *P[gnu-dhd<sup>WT</sup>]* transgene** |  |  |  |
| *dhd<sup>J5</sup> ; P[gnu-dhd]* | *w<sup>1118</sup>* | 331 | 7.25 |
| *dhd<sup>J5</sup> ; P[gnu-dhd]/TM2* | *w<sup>1118</sup>* | 1059 | 3.40 |
| *MTD>sh-lid22 , P[gnu-dhd]/+* | *w<sup>1118</sup>* | 695 | 7.48 |
| **Rescue with *P[UASP-dhd]* inducible transgenes** |  |  |  |
| *nos>+* | *w<sup>1118</sup>* | 711 | 95.36 |
| *nos> P[UASP-dhd]/+* | *w<sup>1118</sup>* | 450 | 97.33 |
| *dhd<sup>J5</sup> ; nos>P[UASP-dhd]* | *w<sup>1118</sup>* | 1163 | 90.28 |
| *dhd<sup>J5</sup> ; nos>P[UASP-dhd<sup>SXXS</sup>]* | *w<sup>1118</sup>* | 702 | 0.00 |
| *MTD>sh-lid22 , P[UASP-dhd]* | *w<sup>1118</sup>* | 1343 | 28.44 |
| *MTD>sh-lid22 , P[UASP-dhd<sup>SXXS</sup>]* | *w<sup>1118</sup>* | 315 | 14.60 |
| *nos>sh-lid21* | *w<sup>1118</sup>* | 1508 | 1.92 |
| *nos> P[UASP-dhd]/sh-lid21* | *w<sup>1118</sup>* | 383 | 19.32 |
| *nos>sh-Sin3a* | *w<sup>1118</sup>* | 542 | 2.03 |
| *nos> P[UASP-dhd]/sh-Sin3a* | *w<sup>1118</sup>* | 621 | 8.86 |
| *nos>sh-Rpd3* | *w<sup>1118</sup>* | 919 | 2.94 |
| *nos> P[UASP-dhd]/sh-Rpd3* | *w<sup>1118</sup>* | 395 | 7.59 |
| *nos>sh-Hira* | *w<sup>1118</sup>* | 514 | 0.00 |
| *nos> P[UASP-dhd]/sh-Hira* | *w<sup>1118</sup>* | 380 | 0.00 |

**Figure 1.**
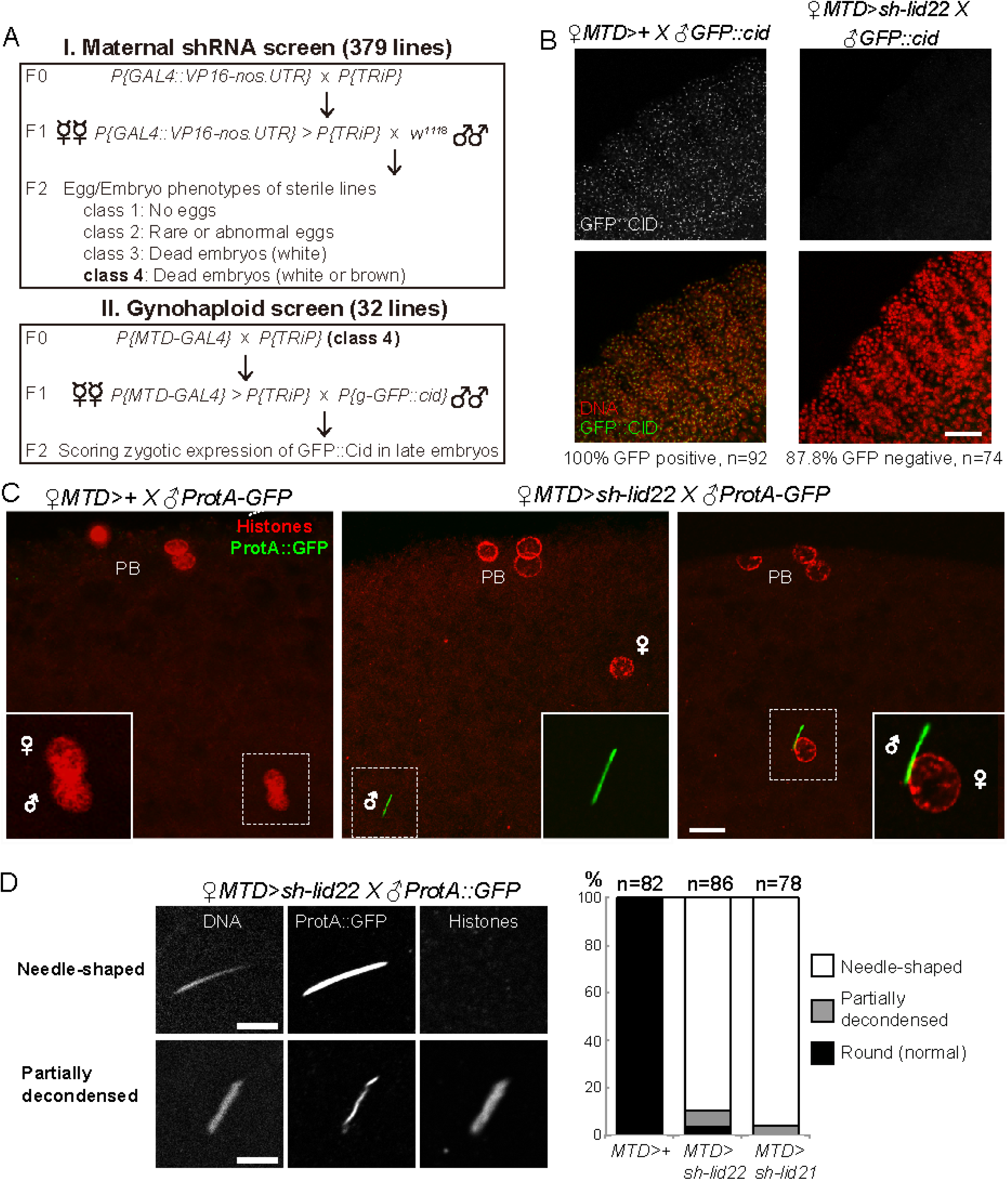
Lid is required for sperm nuclear decompaction at fertilization. A – Scheme of shRNA screen in the female germline. B – Left: Zygotic expression of a paternally-derived *P[g-GFP::cid]* transgene in a control embryo. GFP::Cid marks centromeric chromatin and is visible as nuclear dots. Right: *lid* KD females produce embryos that fail to express the paternally-inherited transgene. Scale bar: 25 µm. C – Maternal Lid is required for SNBP removal and sperm nuclear decompaction at fertilization. Left: a control egg at pronuclear apposition. Both pronuclei (inset) appear similar in size and shape and the SNBP marker ProtA::GFP is not detected. Middle: A representative *lid* KD egg containing a needle-shaped sperm nucleus (inset) still packaged with ProtA::GFP. Right: A fertilized *lid* KD egg with the sperm nucleus apposed to the female pronucleus. Scale bar: 10 µm. D – SNBP replacement with histones is impaired in *lid* KD eggs. Left: Confocal images of representative sperm nuclei in *lid KD* eggs. Partially decondensed nuclei are positive for histones. Scale bar: 5 µm. Right: Quantification of sperm nuclear phenotype in control and *lid* KD eggs.

Two of the identified lines (GLV21071 and GL00612) express the same shRNA against the *little imaginal disc (lid)* gene, which encodes the single fly member of the KDM5/JARID1A family of histone demethylases [10,11]. KDM5 demethylases specifically target the trimethylation of lysine 4 of histone H3 (H3K4me3), a mark typically enriched around the Transcriptional Start Site (TSS) of transcriptionally active genes [12,13]. The two other shRNAs (HMS00359 and HMS00607) target the *Sin3A* and *HDAC1*/*rpd3* genes, respectively. The conserved Sin3A protein scaffold interacts with the histone lysine deacetylase HDAC1 to form the core SIN3 histone deacetylase complex, which is generally considered as a transcriptional repressor [14]. The SIN3 complex regulates the expression of genes involved in a number of metabolic and developmental processes [15–18]. Interestingly, Lid and the largest Sin3A isoform were previously shown to physically and functionally interact [15,16,19,20], thus opening the possibility that these histone modifiers could control the same pathway leading to the formation of a diploid zygote.

### Lid and SIN3 are required for sperm chromatin remodeling at fertilization

The Lid demethylase has been previously shown to be required in the female germline for embryo viability [21,22]. Both studies reported a dual phenotype for eggs produced by *lid* KD females (hereafter called *lid* KD eggs): while a majority of eggs fail to initiate development, a variable but significant fraction developed but died at later stages. Our analysis of *P[g-GFP::cid]* expression (Fig. 1B) indicates that most of these late, non viable KD embryos are haploid and develop with maternal chromosomes. To follow the fate of paternal chromosomes in *lid* KD eggs, we crossed *lid* KD females with males expressing the sperm chromatin marker Mst35Ba::GFP (ProtA::GFP) [23]. In *Drosophila*, protamine-like proteins such as Mst35Ba are rapidly removed from sperm chromatin at fertilization [1] and, accordingly, ProtA::GFP is never observed in the male nucleus of control eggs. In striking contrast, the vast majority of fertilized *lid* KD eggs contained a needle-shaped sperm nucleus that was still positive for ProtA::GFP, indicating that sperm chromatin remodeling was delayed or compromised (Fig. 1C,D). Anti-histone immunostaining indeed revealed that the replacement of SNBPs with maternally supplied histones was variable in KD eggs, ranging from complete absence of histones in the sperm nucleus to the coexistence of variable amounts of histones and ProtA::GFP (Fig. 1D). Notably, we noticed that partially decondensed sperm nuclei in *lid* KD eggs were systematically positive for histones. Taken together, our observations indicate that sperm chromatin remodeling is severely delayed in *lid* KD eggs, thus explaining the absence of paternal chromosomes in most developing embryos. In addition, we observed that, although female meiosis appeared to resume normally in *lid* KD eggs, the female pronucleus frequently (62%, n=76) failed to appose to the sperm nucleus (Fig. 1C). Thus, we propose that defective pronuclear migration accounts for the previously reported failure of *lid* KD embryos to initiate cleavage divisions. Remarkably, we found that germline KD of *Sin3A* also induced a highly penetrant sperm nuclear phenotype with all scored fertilizing sperm nuclei retaining a needle-like shape (100%, n=19) (Fig. EV1). Finally, a similar but less penetrant phenotype was also observed in *rpd3 KD* eggs (Fig. EV1). As this low penetrance could result from less efficient gene knock-down, we chose to mainly focus on *lid and Sin3A* in the rest of our study.

### Transcriptomic analysis of *lid* KD and *Sin3A* KD ovaries identifies *deadhead* as a common and major target gene

In eggs from wild-type females, anti-Lid immunostaining failed to detect Lid protein in the male or female pronucleus, thus suggesting that its implication in sperm chromatin remodeling was indirect. In fact, Lid was not detected in embryos before the blastoderm stage (Fig. EV2). We thus turned to RNA sequencing (RNA Seq) to analyze the respective impact of *lid* KD and *Sin3A* KD on the ovarian transcriptome.

We compared transcriptomes obtained from *lid* KD or *Sin3A* KD ovaries (using the maternal triple driver (MTD-Gal4) with control transcriptomes (MTD-Gal4 only). Globally, our analyses revealed a relatively modest impact of *lid KD* and *Sin3A KD* on ovarian gene expression, with more genes downregulated in both cases (Fig. 2B and EV3A; **Table EV2**). Note that these changes are expected to reflect the activity of Lid and Sin3A in germ cells as ovarian somatic cells (see Fig. 2A) do not express the targeting shRNAs. In their transcriptome analysis (based on microarrays) of *lid* KD wing imaginal discs, Azorin and colleagues found a similar number of differentially-expressed genes, most of them being downregulated [13]. However, RNA Seq analyses recently published by Drelon *et al.* in contrast found 1630 genes (FDR<0.05) dysregulated in wing discs from a null *lid* mutant [24]. Moreover, Liu and Secombe found 8,056 genes differentially expressed (60% were down-regulated) in *lid* adult mutant flies (FDR<0.05), a number which could reflect the greater cell type complexity involved in this analysis [25].

**Figure 2.**
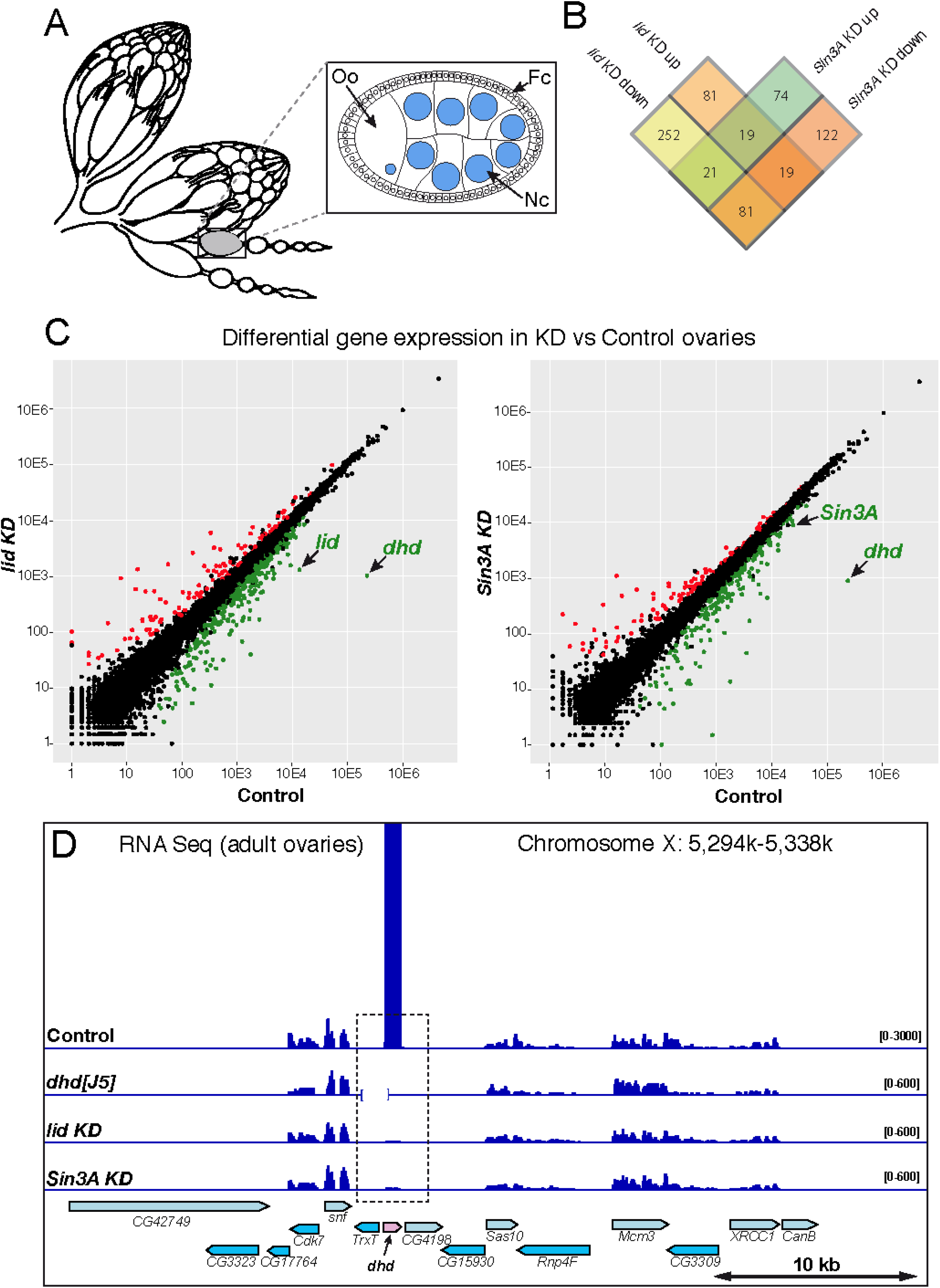
*deadhead* is strongly downregulated in *lid* KD and *Sin3A* KD ovaries. A – Scheme of a pair of adult ovaries with two isolated ovarioles and an egg chamber (inset). Germline nuclei are in blue. Oo: Oocyte, Nc: Nurse cells, Fc: Follicle B – Venn diagram showing the number of differentially expressed genes in *lid* KD and *Sin3A* KD ovarian transcriptomes (FDR<0.001). C – Comparison of RNA seq normalized reads per gene (DESeq2) are shown for *lid* KD vs Control (left) and *Sin3A* KD vs Control (right). Genes with a negative foldchange (downregulated in KD) are in green (FDR<0.001). Genes with a positive foldchange (upregulated in KD) are in red (FDR<0.001). D – Integrative Genomics Viewer (igv) view of Control, *dhd*^*[J5]*^, *lid* KD and *Sin3A* KD ovarian RNA Seq signal on the *dhd* region. The *Df(1)J5* deficiency is indicated as an interrupted baseline on the *dhd*^*[J5]*^ track.

We found that only 29% (139) of the 473 differentially-expressed genes in *lid* KD were also differentially-expressed in *Sin3A* KD, and only 100 genes (21%) were dysregulated in the same direction in both KD (Fig. 2B). As a matter of comparison, Gajan *et al.* found a 65% overlap in *Drosophila* S2 cells [16]. As the *Sin3A* shRNA that we used targets all predicted alternatively spliced mRNAs, it is possible that the knock-down affects Sin3A isoforms with Lid-independent functions [15].

Remarkably, however, we noticed that the *deadhead (dhd)* gene was by far the most severely impacted gene, downregulated by more than two orders of magnitude in both *lid* KD and *Sin3A* KD transcriptomes (Fig. 2C, Fig. EV4). The implication of *dhd* appeared particularly interesting because we and others have recently established that this germline-specific gene is critically required for sperm nuclear decompaction at fertilization [5,26]. *dhd* indeed encodes a specialized thioredoxin that cleaves disufide bonds on SNBPs, thus facilitating their removal from sperm chromatin [5,26]. RT-PCR and Western-blot analyses confirmed the severe repression of *dhd* in *lid* KD and *Sin3A* KD (Fig. EV3).

*dhd* is a small, intronless gene located in the middle of a cluster of fifteen densely packed genes spanning about 40 kb of genomic DNA. Interestingly, the *dhd* gene lies within a 1.4 kb region that is immediately flanked by two testis-specific genes (*Trx-T* and *CG4198*) (Fig. 2D). Despite this apparently unfavorable genomic environment, *dhd* is one of the most highly-expressed genes in ovaries [27], as confirmed by our RNA Seq profiles (Fig. 2C). Interestingly, although this 40 kb region contains six additional genes expressed in ovaries, *dhd* is the only one affected by *lid* KD or *Sin3A* KD (Fig. 2D). Thus, Lid and the SIN3 complex exert a critical and surprisingly specific control on the transcriptional activation of *dhd* in female germ cells.

### Impact of Lid depletion on the distribution of H3K4me3 in ovaries

To evaluate the impact of *lid* KD on the distribution of its target histone mark in the female germline, we performed ChIP-Seq analyses of H3K4me3 in ovaries from control and *lid* KD females. Consistent with earlier reports of a global increase of H3K4me3 in *lid* mutant tissues [13,22,28,29], we observed that H3K4me3 ChIP peaks in *lid* KD ovaries were globally more pronounced compared to control ovaries (Fig. 3A). Our analysis actually revealed that about 10% (1528) of H3K4me3 identified peaks in control ovaries were significantly increased in *lid* KD ovaries (**Table EV3**; FDR<0.05). For those peaks that were associated with genes, the relative enrichment of H3K4me3 in *lid* KD ovaries mainly affected the promoter region and gene body (Fig. 3B). A similar effect was previously observed in *lid* depleted wing imaginal discs, with H3K4me3 abundance specifically increased at the TSS of Lid direct target genes [13]. We nevertheless found 46 H3K4me3 peaks that were significantly decreased in *lid* KD ovaries compared to control ovaries (**Table EV3**; FDR<0.05). Among these, the H3K4me3 peak on the *dhd* gene was the second most severely affected (**Table EV3** and Fig. 3C). Furthermore, only ten of the negatively affected peaks covered genes whose expression were downregulated in *lid KD* ovaries, including *dhd*. Remarkably, the prominent H3K4me3 peak on *dhd* was almost completely lost in *lid* KD ovaries while other peaks within the *dhd* region remained essentially unchanged. *CG4198*, which lies immediately downstream of *dhd* is a notable exception, as this gene also shows a decrease of H3K4me3 despite the fact that it is not expressed in ovaries (Fig. 3C).

**Figure 3.**
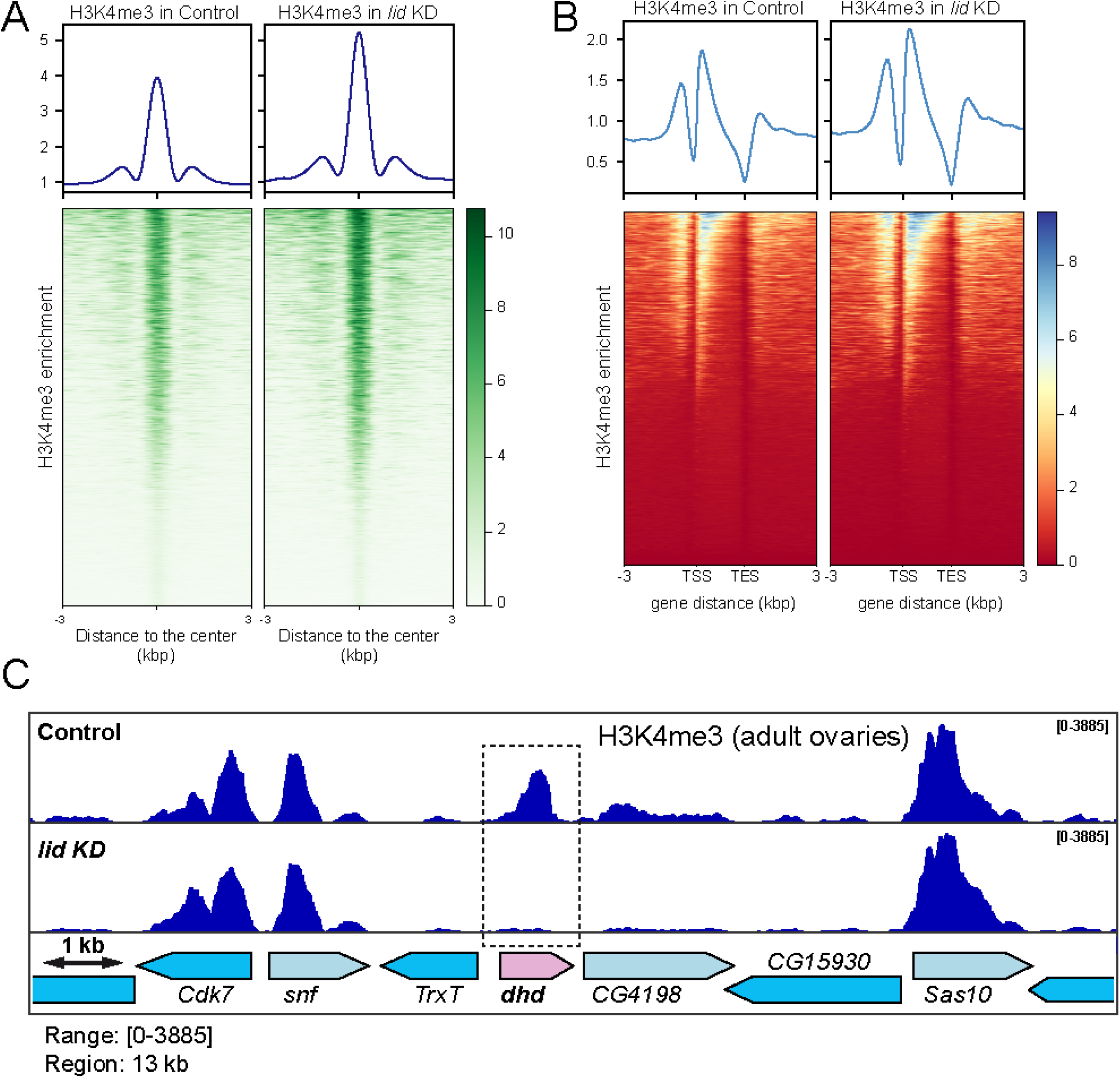
H3K4me3 ovarian ChIP-Seq analysis. A – H3K4me3 enrichment around peak center for Control and *lid* KD ovaries. Upper panels show the average profile around detected peak centers. Lower panels show read density heatmaps around the detected peak centers. B – H3K4me3 enrichment around gene loci for Control and *lid* KD ovaries. Upper panels show the average signal profile on genomic loci defined as 3kb upstream of annotated TSS to 3kb downstream of annotated TES. Lower panels show read density heatmaps around the same genomic loci. C – igv view of H3K4me3 occupancy on the *dhd* genomic region in Control and *lid* KD ovaries.

The paradoxical loss of H3K4me3 enrichment on the *dhd* gene upon Lid depletion is thus another indication that the regulation of this gene is unusual. At least, it indicates that the demethylase activity of Lid is not locally responsible for this regulation. Furthermore, it has been established that *lid* mutant females with a catalytic dead JmjC* *lid* rescue transgene are viable and fertile [24,30]. These rescued females actually have a reduced fertility but still produce about 30% of viable embryos, thus indicating that the demethylase domain of Lid is dispensable to form a viable zygote [22]. KDM5 demethylases can also exert a control on gene transcription by restricting the level of H3K4me3 at enhancer elements [31–33]. The fact that the lysine demethylase activity of Lid is apparently dispensable for *dhd* expression also argues against such a mechanism.

Besides its JmjC demethylase domain, Lid/KDM5 possesses a conserved C-terminal PHD motif capable of binding H3K4me2/3. This binding motif is required for the recruitment of Lid at the promoter of target genes, where it could promote their activation [25]. We thus rather favor a model where the local recruitment of Lid, either through its C-terminal PHD motif or through its DNA binding ARID (AT-rich interaction domain) motif, or both, could establish a chromatin environment permissive to *dhd* massive expression. In this context, the role of the Sin3A/HDAC1 complex also remains to be clarified. The Sin3A histone deacetylase complex is generally considered as a transcriptional repressor [14], but it also functions as a transcriptional activator in *Drosophila* S2 cells [16]. Lacking an intrinsic DNA binding ability [14], the recruitment of this complex to chromatin requires an additional factor. It is thus tempting to propose that Lid itself could recruit Sin3A/HDAC1 locally to activate *dhd* expression in female germ cells. Future elucidation of the mechanisms at play in establishing *dhd* expression will more generally shed light on the role of these yet enigmatic transcriptional regulators.

### Forced *dhd* expression in *lid* KD ovaries partially restores sperm chromatin remodeling at fertilization

Taken together, our cytological and transcriptomic analyses strongly suggest that the loss of *dhd* expression in *lid* KD ovaries at least contributes to the observed fertilization phenotype. To directly test this possibility, we attempted to restore *dhd* expression in *lid* KD female germ cells through the use of transgenic constructs. A genomic transgene (P[*dhd*]) that fully rescued the fertility of *dhd* mutant females [5] only had a very limited impact on the hatching rate of *lid* KD embryos (Table 1) but quantitative RT-PCR analyses revealed that P[*dhd*] remained essentially silent in *lid* KD ovaries (Fig. 4A). This result indicates that the 4.3 kb genomic region present in this transgene is sufficient to recapitulate the endogeneous control exerted by Lid on *dhd* transcription. We then designed another transgene expressing *dhd* under the control of the *giant nuclei* (*gnu*) regulatory sequences. Like *dhd, gnu* is specifically expressed during oogenesis and is functionally required during zygote formation [34]. In addition, our RNA Seq data indicated that its expression is not controlled by Lid. We observed that the *gnu-dhd* transgene indeed restored fertility to *dhd* homozygous mutant females albeit to modest level (about 7% embryo hatching rate; Table 1). In fact, rescued females only produced about 10% of the normal amount of *dhd* mRNA in their ovaries and the DHD protein remained almost undetectable in Western-blot (Fig. 4A). Interestingly, when introduced into *lid* KD females, the *gnu-dhd* transgene also slightly increased embryo hatching rate (Table 1). Furthermore, cytological examination of eggs laid by these females revealed a modest but clear improvement of sperm nuclear decondensation (Fig. 4B,C). These results thus indicate that forced expression of *dhd* can improve the survival of *lid* KD eggs through its positive impact on sperm chromatin remodeling. Finally, we tried to further increase the level of expression of *dhd* by using a Gal4 inducible transgene, P[UAS-*dhd*^*WT*^]. Indeed, induction of this transgene in the germline of *dhd*^*J5*^ mutant females fully restored their fertility (Table 1). We also observed a strong effect on the hatching rate of embryos laid by *lid* KD, P[UAS-*dhd*^*WT*^] females (about 28%; Table 1). However, a P[UAS-*dhd*^*sxxs*^] transgene expressing a catalytic mutant DHD with no rescuing potential that we used as control, also improved the fertility of *lid* KD females, although not as efficiently as the P[UAS-*dhd*^*WT*^] transgene. This effect suggests that Gal4 becomes limiting in the presence of two UAS transgenes, with a negative impact on knock-down efficiency. The fertility of P[UAS-*dhd*^*WT*^] rescued females was nevertheless doubled compared to P[UAS-*dhd*^*sxxs*^] control females, thus supporting the idea that partial *dhd* re-expression in *lid* KD ovaries significantly improved the probability of these eggs to form a viable, diploid zygote. We conclude that the control of *dhd* activation is indeed a critical function of Lid and associated factors in female germ cells.

**Figure 4.**
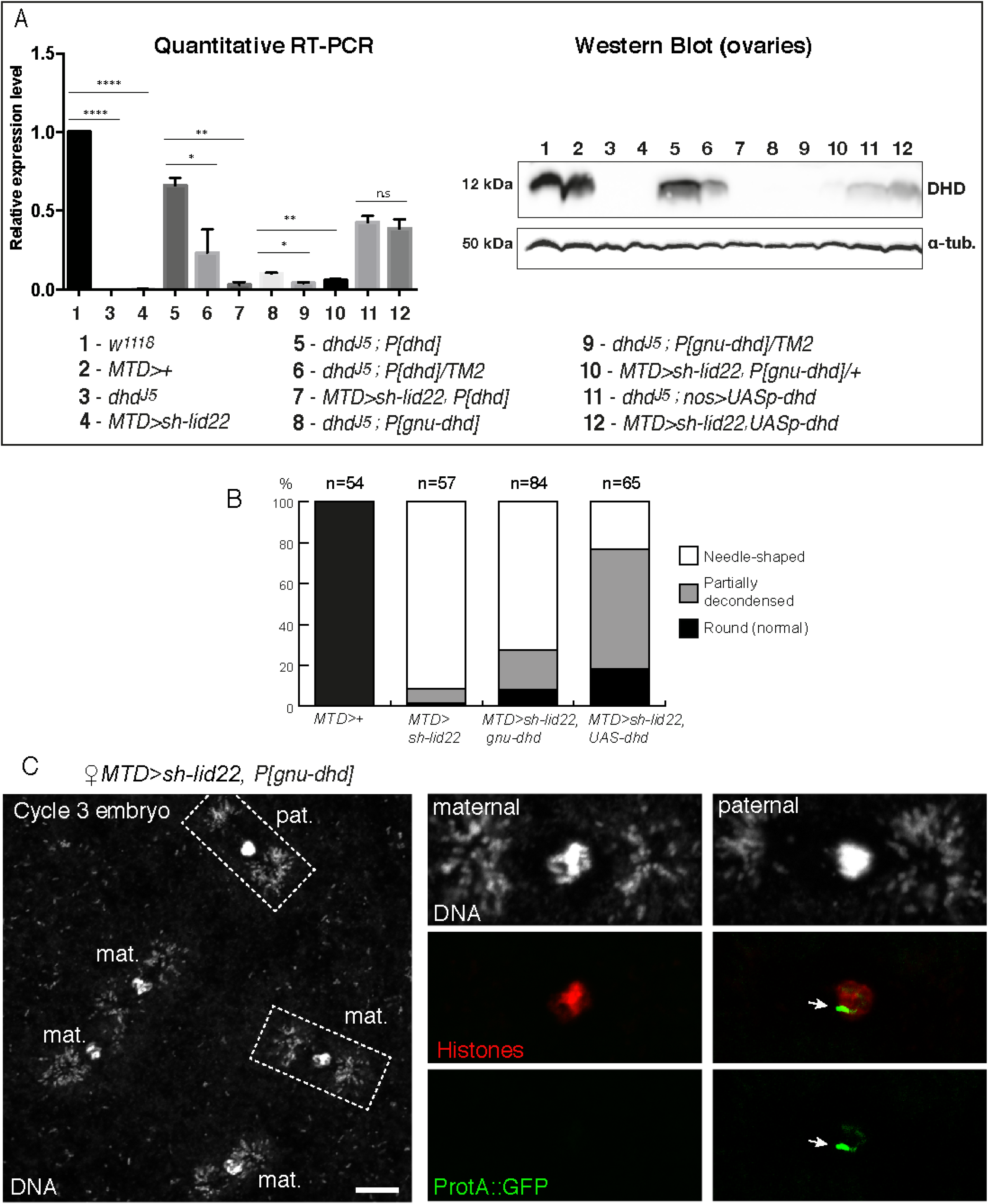
Forced expression of *dhd* partially rescues the *lid* KD phenotype. A – Left: RT-qPCR quantification of *dhd* mRNA levels in ovaries of indicated genotypes (normalized to *rp49* and relative *to* expression in *w*^*1118*^). Data are presented as mean ± SD of 2 biological replicates. P values indicate one-way ANOVA with Dunnett’s multiple comparisons test to a control (**** P < 0.0001; ** P < 0.01, * P < 0.03; n.s = not significant). Right: Western blot analysis of DHD expression in ovaries of indicated genotypes. Alpha-tubulin detection is used as a loading control in Western-blotting. B – Quantification of sperm nuclear phenotype in eggs laid by females of indicated genotypes. C – Confocal images of a *MTD>lid22, P[gnu-dhd]* haploid embryo during the third nuclear division. Karyogamy has failed and the embryo contains four haploid nuclei of presumably maternal origin and one paternal nucleus. The paternal nucleus (inset) still contains a region packaged with ProtA::GFP (arrow). Note that DNA positive dots at the spindle poles are *Wolbachia* endosymbionts. Scale bar: 10 µm.

### Trr controls sperm chromatin remodeling through a Dhd-independent pathway

Intriguingly, germline KD of the Trithorax group protein Trithorax-related (Trr, also known as dMLL3/4), a histone methyltransferase responsible for monomethylation of H3K4 [35], was recently shown to induce a sperm decondensation defect at fertilization similar to the one reported here for *lid*, *Sin3a* and *rpd3* KD [36]. In their study, however, Prudêncio *et al.* did not find any significant change in *dhd* mRNA level in *trr* KD early embryos. To more directly exclude any implication of DHD in the *trr* KD phenotypes, we stained control and KD eggs with an anti-DHD antibody. At fertilization, maternally-expressed DHD is abundant throughout the egg cytoplasm (100%, n=41) but is rapidly degraded after pronuclear apposition. As expected, DHD protein remained undetectable in most *lid* KD eggs (92%, n=50), including those that were fixed before the end of meiosis II (Fig. EV5). In sharp contrast, DHD protein was normally detected in a majority of *trr* KD eggs, even though these eggs indeed contained a needle-shaped sperm nucleus still packaged with SNBPs (Fig. EV5). This result thus confirms that Trr/dMLL3/4 controls sperm nuclear remodeling through a yet unknown, DHD-independent pathway. Conversely, Trr was shown to control meiosis progression through the activation of *Idgf4* [36], a gene that is not affected by *lid* or *Sin3a* KD (this study). Thus Trr and Lid/Sin3A respectively activate a distinct repertoire of genes important for the oocyte-to-zygote transition and sperm chromatin remodeling.

## Conclusion

Our maternal germline genetic screen has unveiled a complex and remarkably specific transcriptional regulation of the *dhd* gene by Lid/KDM5 and the Sin3A/HDAC1 complex. In addition to its crucial role in sperm protamine removal at fertilization, DHD was recently involved in the establishment of a redox state balance at the oocyte-to-zygote transition with a number of identified target proteins [6]. This important DHD-dependent thiol proteome remodeling is thus ultimately controlled by Lid and the SIN3 complex, underlying the critical contribution of these transcriptional regulators to this delicate developmental transition. Future work will aim at dissecting the chromatin mechanisms at play in setting up *dhd* specific activation in female germ cells.

## Materials & Methods

### Drosophila strains

Flies were raised at 25°C on standard medium. The *w*^*1118*^ strain was used as a wild-type control. shRNAs lines used in this study (see **Table EV1**) were established by the Transgenic RNAi Project (TRiP) at Harvard Medical School and were obtained from the Bloomington Drosophila Stock Center at Indiana University. The *lid* and *Sin3A* shRNA lines target all predicted isoforms of their respective target genes. Additional stocks were *EGFP-Cid* [37], *P{otu-GAL4::VP16.R}1; P{GAL4-nos.NGT}40; P{GAL4::VP16-nos.UTR}MVD1* ("MTD-Gal4"), *P{GAL4::VP16-nos.UTR}MVD1* ("nos-Gal4"), *P[Mst35Ba-EGFP]* [23] and *Df(1)J5/FM7c* [38].

### Germline knock-down and fertility tests

To obtain *KD* females, virgin shRNA transgenic females were mass crossed with transgenic Gal4 males at 25°C and females of the desired genotype were recovered in the F1 progeny. To measure fertility, virgin females of different genotypes were aged for 2 days at 25°C in the presence of males and were then allowed to lay eggs on standard medium for 24 hours. Embryos were counted and then let to develop for at least 36 hours at 25°C. Unhatched embryos were counted to determine hatching rates.

### Immunofluorescence and imaging

Early (0-30 min) and late (about 6 hours) embryos laid by randomly selected females were collected on agar plates. Embryos were dechorionated in bleach, fixed in a 1:1 heptane:methanol mixture and stored at −20°C. Embryos were washed three times (10 min each) with PBS1X 0.1%, Triton X-100 and were then incubated with primary antibodies in the same buffer on a wheel overnight at 4°C. They were then washed three times (20 min each) with PBS 0.1%, Triton X-100. Incubations with secondary antibodies were performed identically. Embryos were mounted in Dako mounting medium containing DAPI.

Ovaries were dissected in PBS-Triton 0.1% and fixed at room temperature in 4% formaldehyde in PBS for 25 minutes. Immunofluorescence was performed as for embryos except for secondary antibodies that were incubated four hours at room temperature. Ovaries were then mounted as described above.

Primary antibodies used were mouse monoclonal anti-histones (Sigma #MABE71; 1:1000), rabbit polyclonal anti-DHD (1:1000) [5], rat polyclonal anti-Lid (1:500) [13] and mouse monoclonal anti-GFP (Roche #118144600001; 1:200). Secondary antibodies were goat anti-rabbit antibodies (ThermoFisher Scientific, 1:500) and goat anti-mouse antibodies (Jackson ImmunoResearch, 1:500) conjugated to AlexaFluor. Images were acquired on an LSM 800 confocal microscope (Carl Zeiss). Images were processed with Zen imaging software (Carl Zeiss) and Photoshop (Adobe).

### Western Blotting

Ovaries from 30 females were collected and homogenized in lysis buffer (20mM Hepes pH7.9, 100mM KCl, 0.1mM EDTA, 0.1mM EGTA, 5% Glycerol, 0.05% Igepal and protease inhibitors (Roche)). The protein extracts were cleared by centrifugation and stored at −80°C. Eggs were collected every 30 min, dechorionated in bleach and quickly frozen in liquid nitrogen. Protein extracts were prepared from ca. 10 µl of embryos. Protein samples were run on 15% SDS polyacrylamide gel and transferred to Immun-Blot^®^ PVDF membrane (Bio-Rad) for 1h at 60V. Membranes were blocked for 1h at room temperature in 5% non-fat milk in PBS 1X-Tween20 0.05%, followed by an overnight incubation with the primary antibody at 4°C in 5% non-fat milk in PBS1X-Tween20 0.05%. Secondary antibodies used were added and incubated for 2 hours at room temperature. Protein detection was performed using ECL solution according manufacturer’s instruction (GE Healthcare). Antibodies used were: rabbit polyclonal anti-DHD (1/1000) [5], mouse monoclonal anti-α-Tubulin (Sigma Aldrich #T9026, 1:500), HRP-conjugated goat anti-mouse (Biorad #170-5047; 1:50 000) and peroxidase-conjugated goat anti-rabbit (Thermoscientific #32460; 1:20 000).

### Gene expression analysis by RT-QPCR

Total RNA was extracted from ovaries of 3-day-old females using the NucleoSpin® RNA isolation kit (Macherey-Nagel), following the instructions of the manufacturer. Duplicates were processed for each genotype. cDNAs were generated from 1µg of purified RNA with oligo (dT) primers using the SuperScript^TM^ II Reverse Trancriptase kit (Invitrogen).

Generated cDNAs were diluted to 1/5 and were used as template in a real time quantitative PCR assay using SYBR®Premix Ex TaqTM II (Tli RNaseH Plus) (Takara). All qRT-PCR reactions were performed in duplicate using Bio-Rad CFX-96 Connect system with the following conditions: 95°C for 1 min followed by 40 cycles of denaturation at 95°C for 10 s, annealing at 59°C for 30 s and extension at 72°C for 30 s. Relative fold change in gene expression was determined by the comparative quantification ΔΔCT method of analysis [39]. The housekeeping gene *rp49* was used to normalize cDNA amounts in the comparative analysis. The primer sets used in the PCR reactions were: dhd-forward 5’-TCTATGCGACATGGTGTGGT −3’ and dhd-reverse 5’-TCCACATCGATCTTGAGCAC −3’; Rp49-forward 5’-AAGATCGTGAAGAAGCGCAC −3’ and Rp49-reverse 5’-GATACTGTCCCTTGAAGCGG −3’. Statistical tests were performed using GraphPad Prism version 6.00 for Mac OS X (GraphPad Software).

### Ovarian RNA Sequencing and analysis

For each samples, 8 pairs of ovaries were dissected from 6 day-old virgin females and total RNA were extracted using the NucleoSpin® RNA isolation kit (Macherey-Nagel), following the instructions of the manufacturer. Extracted RNAs were treated with Turbo™DNAse (Ambion #AM2238). After DNase inactivation, RNAs were purified using the NucleoSpin® RNA Clean-up XS kit (Macherey-Nagel) according to manufacturer’s instructions. Sequencing was completed on two biological replicates of each genotype:

Control KD (MTD-Gal4>+)

P{w[+mC]=otu-GAL4::VP16.R}1, w[*]/y[1] v[1]; P{w[+mC]=GAL4-nos.NGT}40/+;

P{w[+mC]=GAL4::VP16-nos.UTR}CG6325[MVD1]/P{y[+t7.7]=CaryP}attP2

lid KD (MTD-Gal4>shRNA lid)

P{w[+mC]=otu-GAL4::VP16.R}1, w[*]/y[1] sc[*] v[1]; P{w[+mC]=GAL4-nos.NGT}40/+;

P{w[+mC]=GAL4::VP16-nos.UTR}CG6325[MVD1]/P{y[+t7.7] v[+t1.8]=TRiP.GLV21071}attP2

lid KD (MTD-Gal4>shRNA Sin3A)

P{w[+mC]=otu-GAL4::VP16.R}1, w[*]/y[1] sc[*] v[1];P{w[+mC]=GAL4-nos.NGT}40/+;

P{w[+mC]=GAL4::VP16-nos.UTR}CG6325[MVD1]/P{y[+t7.7] v[+t1.8]=TRiP.HMS00359}attP2

Sequencing libraries for each sample were synthesized using TruSeq Stranded mRNA kit (Illumina) following supplier recommendations (Sample Preparation Guide - PN 15031047, version Rev.E Oct 2013) and were sequenced on Illumina Hiseq 4000 sequencer as Single-Reads 50 base reads following Illumina’s instructions (GenomEast platform, IGBM, Strasbourg, France). Image analysis and base calling were performed using RTA 2.7.3 and bcl2fastq 2.17.1.14. Adapter dimer reads were removed using DimerRemover. Sequenced reads were mapped to the *Drosophila melanogaster* genome assembly dm6 using TopHat (version 2.1.1) with default option. The aligned reads were assigned to genes by FeatureCounts, run with default options on the dmel-all-r6.15 version of the *Drosophila melanogaster* genome annotation. Differentially expressed genes were identified using the R-package DESeq2 (version 1.14.1). The annotated genes exhibiting an adjusted-P < 0.001 were considered differentially expressed compared to Control.

### Chromatin immunoprecipitation, sequencing and analysis

ChIP assays were performed as previously described [40]. Two biological replicates for *control* KD and *lid* KD ovaries (same genotypes as for RNA Seq) were processed and analyzed. For each biological replicate, eighty ovary pairs were dissected from 2 day-old females and flash frozen. Dissected ovaries were fixed in 1.8% formaldehyde at room temperature for 10 minutes. Chromatin was sonicated using a Diagenod Bioruptor (18 cycles, high intensity, 30s on/30s off) to generate random DNA fragments from 100 to 800 base pairs. Sheared chromatin was incubated overnight at 4°C with H3K4me3 antibody (ab8580 Abcam). Immunoprecipitated samples were treated with RNAse A, proteinase K and DNA purified using the ChIP DNA Purification kit (Active Motif #58002) following the manufacturer’s instructions. Quantification assessment of purified DNA was done using Qbit dsDNA HS Assay on the Qbit fluorometer (Invitrogen). Immunoprecipited DNA quality was evaluated on a Bioanalyzer 2100 (Agilent).

Sequencing libraries for each sample were synthesized using Diagenode MicroPlex Library Preparation kit according to supplier recommendations (version 2.02.15) and were sequenced on Illumina Hiseq 4000 sequencer as Paired-End 50 base reads following Illumina’s instructions (GenomEast platform, IGBM, Strasbourg, France). Image analysis and base calling were performed using RTA 2.7.3 and bcl2fastq 2.17.1.14. Adapter dimer reads were removed using DimerRemover. Sequenced reads were mapped to the *Drosophila melanogaster* genome assembly dm6 using Bowtie (version 2.3.3) with default option. Only uniquely aligned reads have been retained for further analyses. Duplicated reads were removed using picard-tools (version 2.17.10). Peak calling was performed for each individual samples and on merged biological replicates using MACS algorithm (version 2.1.1) with default option and a relaxed q-value cut-off of 0.1. Consistent peaks between biological replicates were identified using irreproducible discovery rate (IDR version 2.0.3) with a 0.05 cut-off. Differentially modified H3K4me3 peaks between Control and *lid* Knock-down ovaries were identified using the R-package DiffBind (version 2.2.12) with a 0.05 FDR cut-off.

### Data visualization

The Deeptools software was used to convert alignment files to bigwig (bamCoverage) and to generate H3K4me3 heatmap and density profiles (computeMatrix and plotHeatmap). The generated bigwig files were visualized using IGV software.

## Data availability

The DNA sequencing data from this publication have been deposited to the Gene Expression Omnibus database [https://www.ncbi.nlm.nih.gov/geo/] and assigned the identifier GSE133064 (RNA-Seq) and GSE133202 (ChIP-Seq).

## Acknowledgments

We thank Cristina Molnar, Cayetano Gonzalez and Ferran Azorin for sharing fly stocks and antibodies. We are grateful to the TRiP projects and Bloomington Drosophila Stock Center for shRNA stocks. The authors would like to thank the members of the IGBMC GenomEast platform, member of the "France Génomique" consortium (ANR-10-INBS-0009). We also thank Raphaëlle Dubruille for her helpful comments on the manuscript. SK was supported by a fellowship from the ANR (ZygoPat ANR-12-BSV6-0014). DT is supported by a doctoral fellowship from the French government.

---

**Figure EV1.**
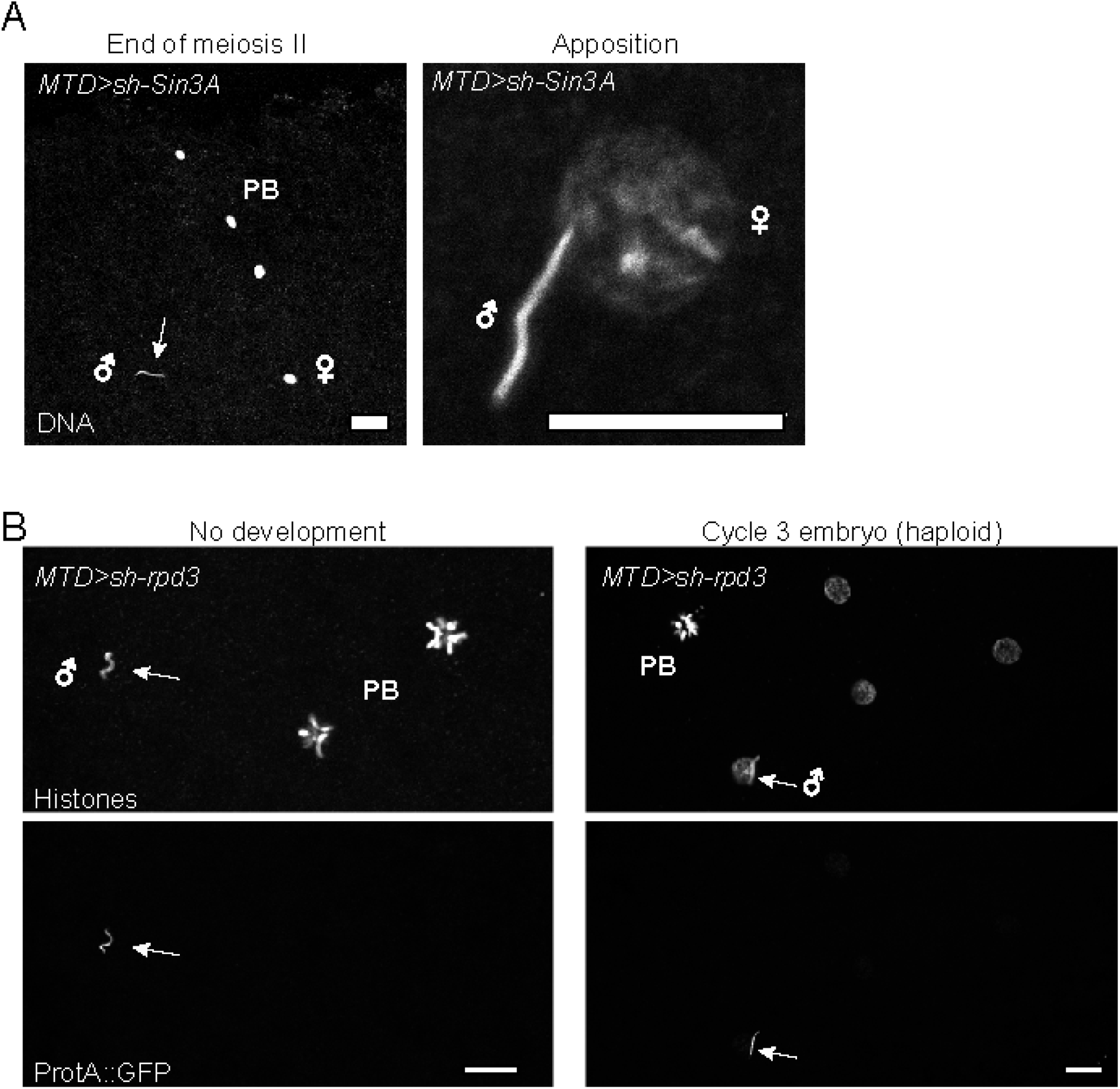
Phenotype of *Sin3A* KD and *rpd3* KD e1:1:s/em bryos. A – Confocal images of *Sin3A* KD eggs stained for DNA at the indicated stages. The sperm nu cleus in the left panel is indicated (arrow). Bar: 10 µm PB: Polar bodies. B – Confocal images of *rpd.3* KD early emb_ry_os (from ProtA::GFP fathers) stained for DNA and anti-GFP The sperm nu cleus is indicated (arrows). Bar: 10 *µm.* PB: Polar bodies.

**Figure EV2.**
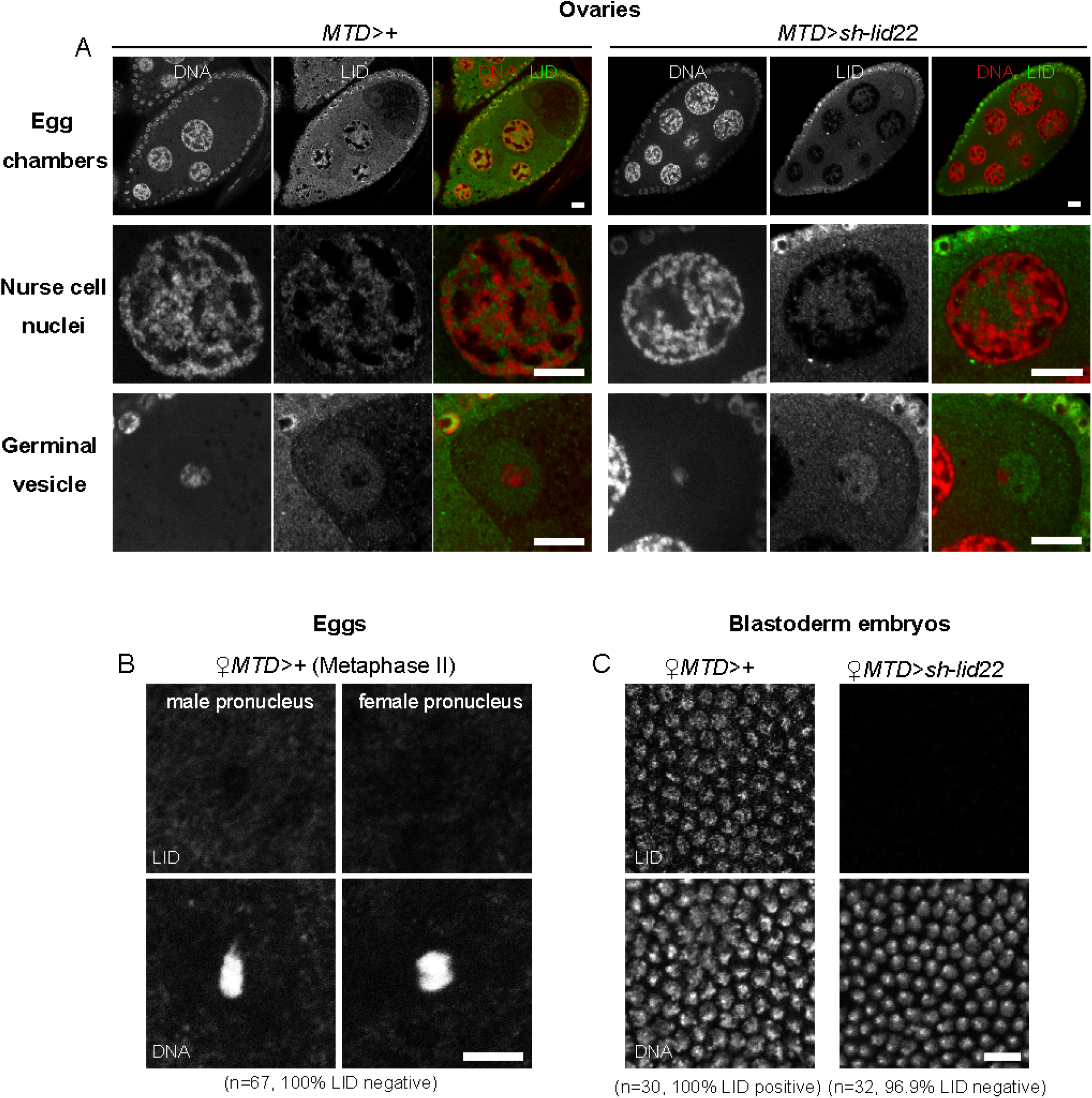
Lid is not directly invoved in sperm chromatin remodelinz:: at fertilization. A – Top row: confocal images of stage 10 egg chambers from control (left) and *lid* KD (right) females stained for DNA (red) and anti-Lid (green). Middle row: detail of a nurse cell nucleus. Bottom row: detail of the oocyte germinal vesicle (oocyte nucleus). Bar: 20 *µm*. B – Confocal images of the male pronucleus and the female pronucleus from a control egg in meiosis II stained for DNA and anti-Lid. Bar: 10*µ*m Quantification of Lid positive nuclei is indicated. C – Confocal images of a control (left) and *lid* KD (right) blastoderm embryo with same stainings as in B. Bar: 10 *µm.* Quantifications of embryos with a positive/negative nuclear Lid staining are indicated for each genotype.

**Figure EV3.**
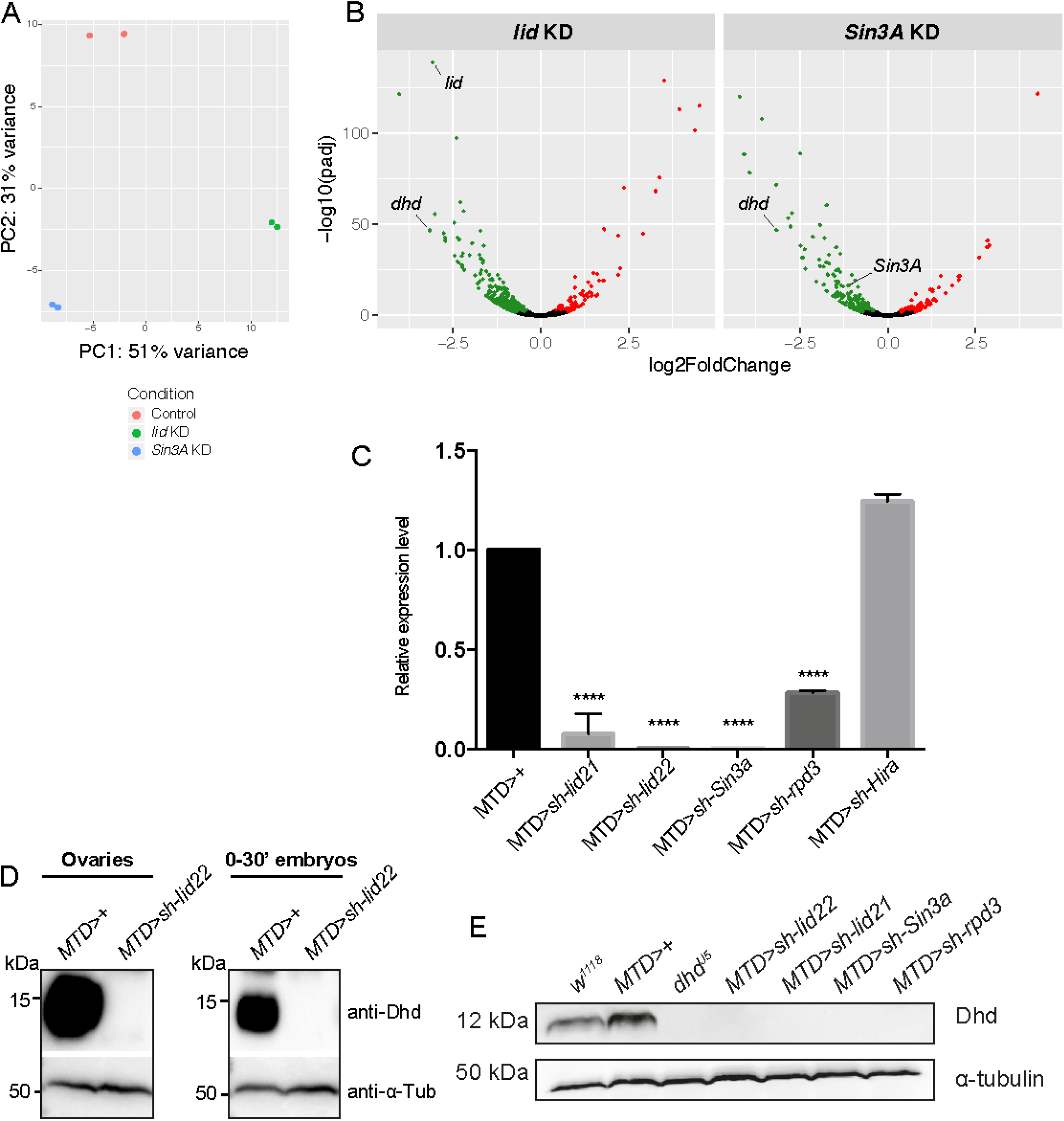
Lid and Sin3A control dhd expression in female a=erm cells. A – Principal Component Analysis of Control, *lid* KD and *Sin3A* KD ovarian transcriptomes (two biological replicates for each genotype). B – Volcano plot representations of Differentially-Expressed genes in Control vs *lid* KD (left) and Control vs *Sin3A* KD (right). C – RT-qPCR quantification of *dhd* mRNA levels in ovaries of indicated genotypes. mRNA levels were normalized to *rp49* and shown as relative expression in MTD>+ control. Error bars represent SD (Dunnett’s multiple comparisons test to the control MTD>+, **** P < 0.0001). D – Western blot analysis of DHD in adult ovaries (left) and 0-30min post fertilization embryos (right). a-tubulin was used as a loading control. E – Western blot analysis of DHD in adult ovaries of indicated genotypes. a-tubulin was used as a loading control.

**Figure EV4.**
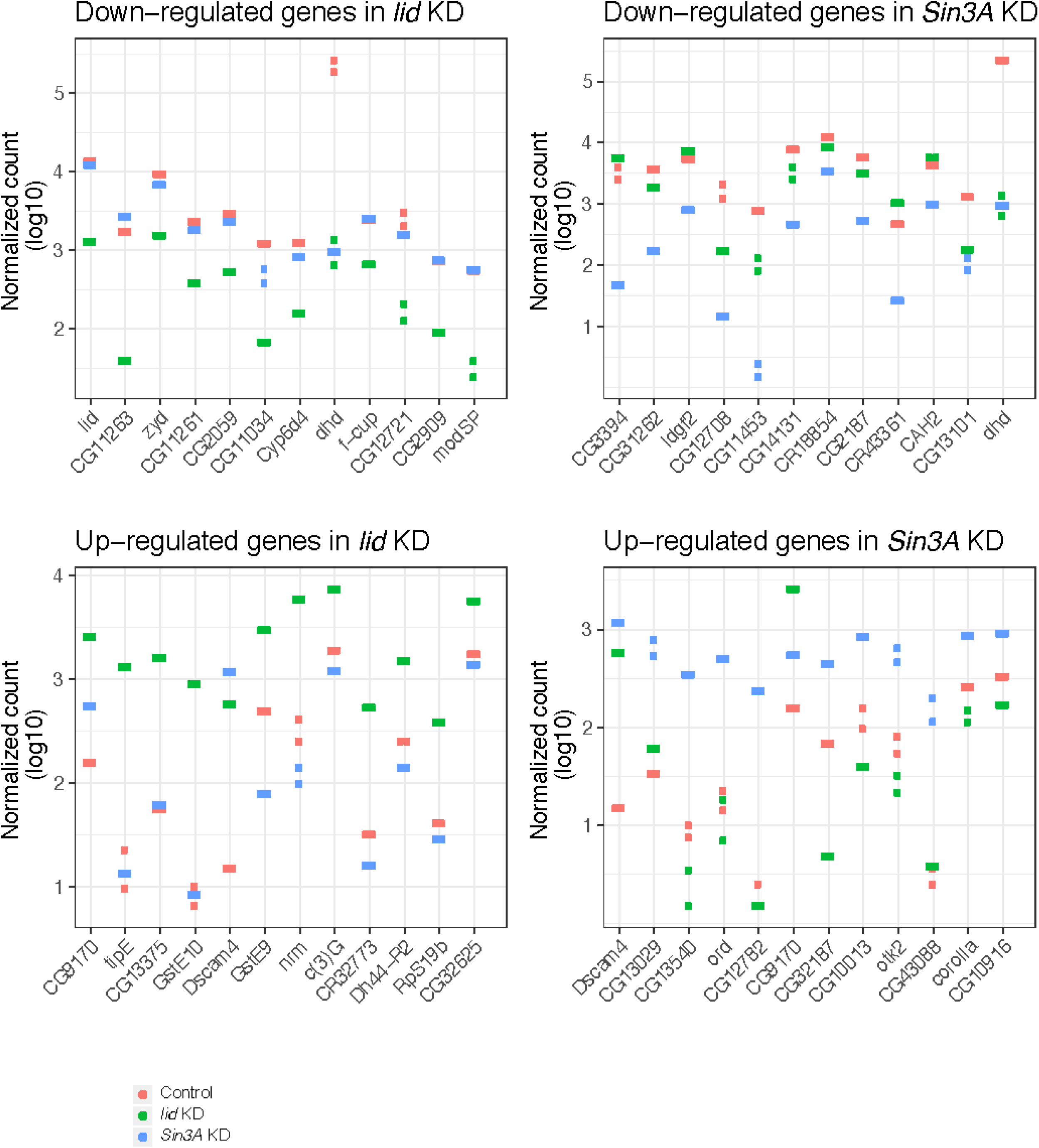
Top twelve most downregulated and upregulated genes in *lid* KD and *Sin3A* KD transcriptomes.

**Figure EV5.**
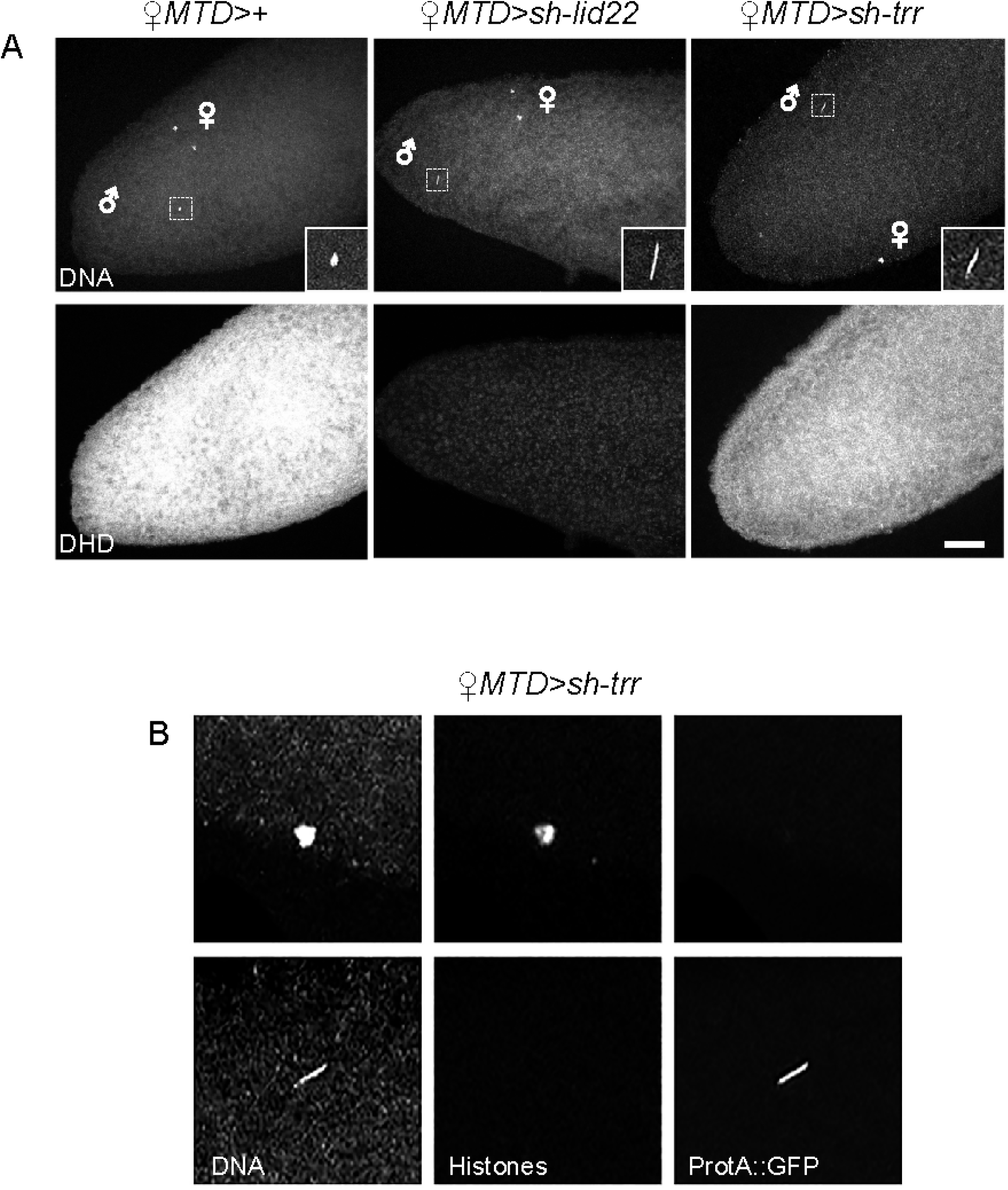
*trr* KD does not affect *dhd* expression. A – Confocal images of representative embryos of the indicated genotypes stained for DNA and anti-DHD. The fertilizing sperm nucleus is magnified in insets. Bar: 20*µ*m B – Details of maternal chromosomes (top row) and sperm nucleus (bottow row) from a representative *trr* KD egg stained for ProtA::GFP and histones.

